# A family portrait of lanmodulin selectivity for enhanced rare-earth separations

**DOI:** 10.64898/2026.01.22.699517

**Authors:** Patrick Diep, Cody S. Madsen, Wonseok Choi, Ziye Dong, Christina S. Kang-Yun, Patricia F.V. Uychoco, Jeremy A. Seidel, Samuel A. Eaton, Yongqin Jiao, Joseph A. Cotruvo, Dan M. Park

## Abstract

Proteins offer a molecular design space to create bespoke ligands for the separation of critical metals, like rare earth elements (REs). However, data-intensive approaches to fine-tune metalloprotein selectivity are bottlenecked by the low-throughput nature of existing characterization methods. Here we invented an assay called *SpyTag-Catcher Immobilization of Lanmodulin for Assaying Metal-Binding Selectivity* (SpyCI-LAMBS) to measure metalloprotein selectivity en masse. This 96-format workflow was used to study the selectivity of 621 lanmodulin (LanM) orthologs for 15 REs, revealing eight distinct selectivity profiles based on sequence-to-function analyses. We discovered >200 LanMs with dampened selectivity for low-value La^III^ relative to the prototypical LanM. This includes a LanM that can perform a challenging one-stage separation of Pr^III^ from La^III^ with up to >99.9 mol% purity and 83% yield. SpyCI-LAMBS is a powerful tool that can rapidly collect high-fidelity selectivity data to inform metal ion separations and machine learning-assisted metalloprotein design.

## Introduction

Global supply chain instability, heightened since the COVID-19 pandemic and exacerbated by international geopolitical tensions, persists alongside the pressing need to strengthen climate resilience through increased deployment of clean energy technologies that require rare earth elements (REs) and other strategic metals.^1,2^ Due to the environmental impacts of traditional mining practices, new economical approaches are needed to diversify supply chains to sustain technological advancements. The production of highly pure individual REs is especially difficult due to their similar physicochemical properties (*i.e*., ionic radius, coordination number and geometry) and natural co-occurrence^3^. At present, the main approach to purify individual REs is through multi-stage liquid-liquid extraction^4^. To avoid the immense volumes of organic solvents and acid required in this conventional approach, new protein-based biotechnologies for all-aqueous RE extraction and purification are being developed^5–16^.

The lanthanide binding protein lanmodulin (LanM) from *Methylobacterium extorquens* (*Mex-*LanM) has emerged as a promising candidate for RE purification. *Mex*-LanM exhibits extraordinarily high selectivity for REs^5,6,17^ and sufficient intra-RE selectivity when immobilized in column format to achieve individual separation between some RE pairs (*e.g*., Nd^III^ vs. Dy^III^)^6,11^. However, *Mex-*LanM exhibits similar affinity for Ce^III^ through Eu^III^, which limits the separation potential between light REs (LRE; La^III^-Sm^III^)^11,17,18^. A recent effort identified *Hans-*LanM from *Hansschlegelia quercus* in a bioinformatic campaign that unearthed 696 putative LanM orthologs^19^. *Hans*-LanM exhibits lanthanide-dependent dimerization and substantially enhanced selectivity for La^III^ to Nd^III^ compared to heavy REs (HREs; Eu^III^-Lu^III^, Y^III^). When immobilized, *Hans*-LanM enabled single-stage separation of a 95:5 Nd^III^/Dy^III^ solution^19^.

LanM’s RE selectivity arises from an intricate interplay between local (*e.g*., metal-binding EF-hands) and global (*e.g*., long-range interactions in the fully folded state) structure^20,21^. For example, *Hans*-LanM’s enhanced selectivity is thought to arise from an exclusively protein-derived first coordination sphere modulated by a hydrogen-bonding network at the dimer interface^19^. Conversely in *Mex*-LanM, hydrogen bonds between two coordinating water molecules and second-sphere residues are critical for its metal binding function^20,22^. The distinct coordination strategy and RE selectivity between *Hans*-LanM and *Mex*-LanM highlight the potential diversity that exists among the natural LanM family, motivating further experimental characterization of LanM orthologs.

To engineer LanMs to selectively bind the more valuable HREs or other critical metals^23^, a broader understanding of their sequence-structure-selectivity relationship is needed. Machine learning paradigms for protein design may generate new LanMs with bespoke selectivity profiles, but this approach requires larger and more diverse datasets for model training. Herein lies the major bottleneck for using selectivity as a design objective – it incurs significant resources to experimentally quantify, yet it is the most direct predictor of separation performance. In metal separations, the commonly used metric to quantify a ligand’s selectivity is the distribution coefficient (*D*). In prior work, we developed a column-based approach to determine LanM’s *D* values for each RE present in a total RE (“tRE”: Y^III^, La^III^-Lu^III^, except Pm^III^) equimolar solution, partitioned into the solid phase (LanM) and the aqueous phase (the unbound fraction). At least a week is ideally needed to quantify selectivity (*D*) for at most a handful of proteins with this column-based approach, from protein production to column testing and analysis (Supplementary Table 1). Consequently, exploring LanM’s natural selectivity landscape has been untenable for larger libraries (10^2^-10^3^), thus precluding extensive engineering campaigns and a deeper exploration of the natural diversity of selectivities within this family. Higher throughput methods to measure the selectivity of LanM for REs, and in general any metalloprotein for any set of metals, are necessary to expand the palette of proteins for metal separations.

Here, we report the invention of the SpyCI-LAMBS assay (*<u>Spy</u>Tag-<u>C</u>atcher Immobilization of <u>L</u>anmodulin for <u>A</u>ssaying <u>M</u>etal-<u>B</u>inding <u>S</u>electivity*). SpyCI-LAMBS enables expression, immobilization, and quantification of selectivity for hundreds of proteins within a week (not including cloning). We demonstrated the utility of SpyCI-LAMBS by measuring the intra-RE selectivity of 621 LanM orthologs to assess the breadth of natural selectivity profiles^19^, which revealed a portrait of eight distinct selectivity clusters (herein “clusters” for brevity), including a large cluster with La^III^ rejecting behavior not previously reported. We then translated the principles of SpyCI-LAMBS to column format in a method we named SpECS (*<u>S</u>pyCI-LAMBS for <u>E</u>stimating <u>C</u>hromatographic <u>S</u>eparations*) to enable screening of separation potential for a panel of orthologs using high aspect ratio columns. This identified a LanM ortholog from *Methylobacterium* sp.13MFTsu3.1M2 (called “o-36”) that achieved single-stage separation of the near-adjacent Pr^III^/La^III^ pair, a challenging separation for *Mex*-LanM and *Hans*-LanM. These results illustrate the power of SpyCI-LAMBS to decrease the turnaround time for measuring the selectivity of metalloproteins, which is needed to predict their separation potential and to feed machine learning models for metalloprotein design.

## Results

### Quantifying selectivity by SpyCI-LAMBS

Our goal was to miniaturize the column-based approach for measuring the RE selectivity of LanMs in 96-well format. We began in microfuge tube format by first deploying SpyTag-Catcher conjugation chemistry^24^ to directly immobilize SpyTagged LanM protein from lysate and thus avoid all protein purification steps (Figure 1a), similar to what was previously demonstrated with magnetic nanoparticles^8^. Controlled pore glass (CPG) was chosen as the solid matrix resin due to its inertness and high density relative to water, which enabled facile liquid-solid separation of bound REs from the aqueous phase. As a proof-of-concept, we prepared a lysate containing SpyTagged super folder (sf)GFP and conjugated it to CPG functionalized with SpyCatcher (herein “CPG:SpyCatcher”). We observed attachment of sfGFP based on fluorescence microscopy (Figure 1b) and a 96% reduction in lysate fluorescence (Supplementary Figure 1). Using an ascending gradient of CPG:SpyCatcher slurry, but keeping the lysate volume constant, we then confirmed removal of SpyTagged *Mex-*LanM from lysate in a proportional manner (Figure 1c). These results established the SpyTag-Catcher pair could enable specific attachment of target proteins to resin directly from lysate.

**Figure 1.**
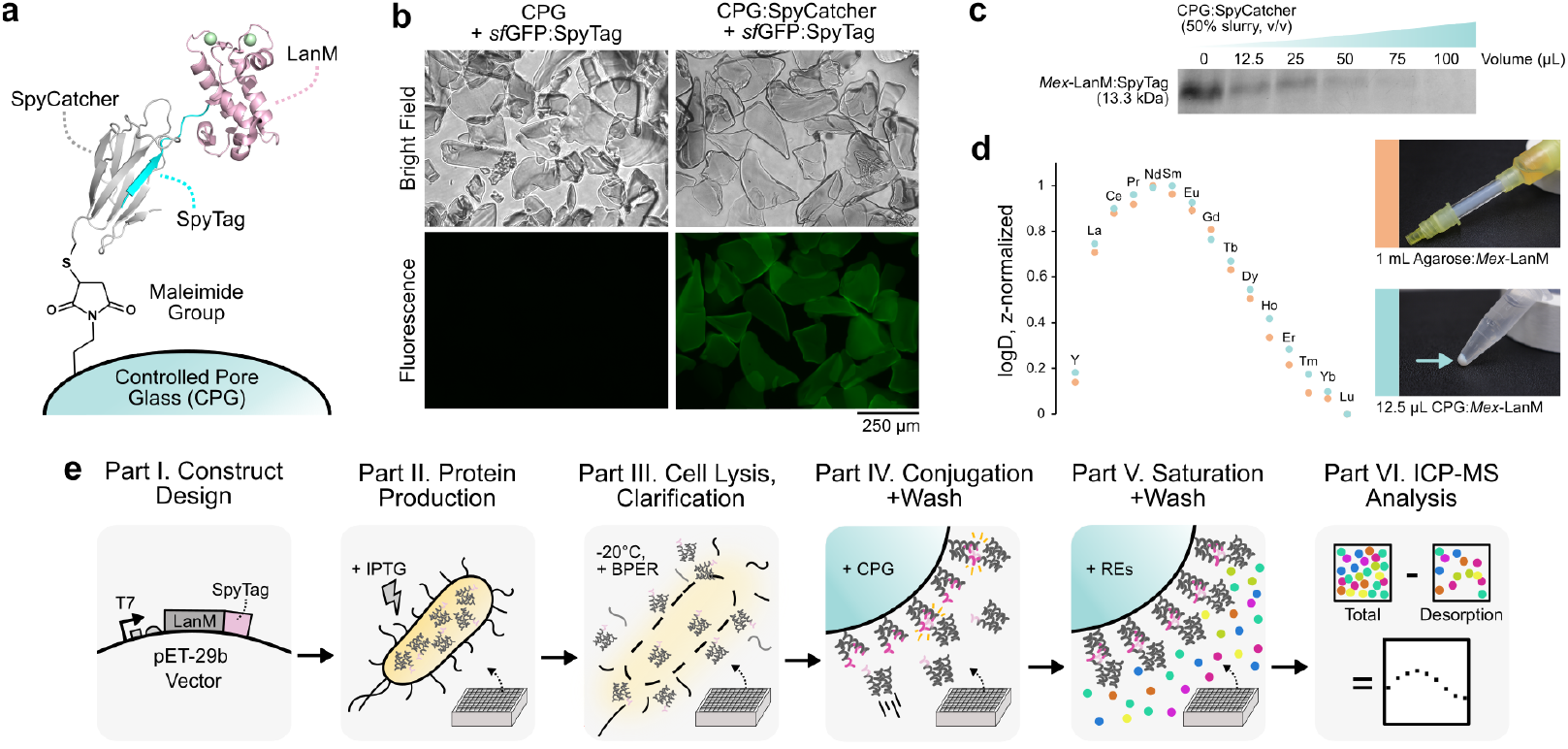
SpyCI-LAMBS overview. (a) Concept. The first component is “CPG:SpyCatcher,” a slurry of SpyCatcher immobilized to porous glass particles (CPG), chosen due to its innertness and high density relative to water that enables facile liquid-solid separation. The second component is “LanM:SpyTag,” a clarified lysate of soluble LanM protein with SpyTag3 appended to the C-terminus. The 50% (v/v) slurry product of the two conjugated components is referred to as “CPG:LanM”. (b) Visual confirmation of SpyTag-Catcher immobilization of sfGFP:SpyTag to CPG:SpyCatcher by fluorescence microscopy. n =1. (c) SDS-PAGE confirmation of LanM:SpyTag removal from lysate by immobilization to CPG:SpyCatcher. n = 1. (d) log*D* congruency for *Mex*-LanM between the column-based approach (orange) and a SpyCI-LAMBS proof-of-concept (blue). n = 1. (e) The 96-deepwell SpyCI-LAMBS workflow has six parts: Part I) cloning or DNA synthesis to produce the T7 expression vector harboring the LanM ortholog or variant of interest, Part II) protein expression with IPTG induction, Part III) lysis and clarification by centrifugation to obtain a clean soluble fraction, Part IV) addition of CPG:SpyCatcher to the soluble fraction for immobilization of the target LanM:SpyTag, Part V) a mixture of metals, such as a total RE (tRE) solution, is added to the CPG:LanM at a pre-determined ratio for equilibration, and Part VI) after washing, the metal is desorbed and analyzed by ICP-MS to compute the log*D* values from the determined concentrations.

To tune SpyCI-LAMBS for the largest dynamic range, two parameters were varied: 1) the resin quantity and 2) the metal concentration (Extended Figure 1, Supplementary Figure 2, Supplementary Figure 3, Supplementary Note 1). We determined 25 μL of a 50% (v/v) CPG:*Mex*-LanM slurry saturated with 1 mL of equimolar tRE solution (each RE at 100 ppb, buffered at pH 3.0) had the largest signal-to-noise ratio. Next, we computed log*D* values for each metal in the tRE solution (Supplementary Note 2) and compared this selectivity profile with that determined using the column-based approach (Figure 1d, Extended Figure 2a,b). These experiments confirmed that *Mex-*LanM assayed for selectivity in small-volume SpyCI-LAMBS format (Figure 1e) could produce the same log*D* profile as our established, but lower throughput, column-based approach.

To assess SpyCI-LAMBS’ reproducibility and to optimize the workflow to generate consistent log*D* profiles for CPG:*Mex*-LanM, we tested the material’s behavior by varying eight parameters (Extended Figure 3). Of the parameters assessed, only the serial dilution of the lysate, meant to mimic low-expression proteins, led to a marked flattening of the log*D* profile (Extended Figure 3g). We found these changes in log*D* values to be non-linear, and that 10% and even 1% of *Mex*-LanM’s expression level could still generate useful selectivity trends. In other words, log*D* profiles are robust to differences in soluble protein expression levels over a nearly 100-fold range, which is important for protein variants with weak protein expression not known *a priori*. Nearly identical log*D* profiles were observed when five consecutive replicates with the same preparation of CPG:*Mex*-LanM material were performed, as well as when the material was stored at 4 °C in water for up to a month (Extended Figure 2c-g). These optimized parameters were then translated to 96-deepwell block format, which we confirmed had negligible cross-contamination between wells (Supplementary Figure 4-Supplementary Figure 5)^25^. Altogether, this work establishes a highly sensitive, reproducible, and robust assay to search for new RE selectivities in both tube and block format.

### Snapshots of the LanM selectivity landscape

In our prior work, 696 putative LanM orthologs were identified using a PSI-BLAST against the NCBI database and filtering for sequences with <200 residues, 4 EF-hands in total, and at least 1 pair of EF-hands separated by <14 residues^19^. From this set, we extracted a subset of 621 orthologs (herein termed the “library”) after manually removing redundant 4 EF-hand sequences and re-adding 3 EF-hand orthologs (Supplementary Data 1). The identity of each ortholog is denoted as “o-X”, where “X” demarks the list position indexed at 0 (*e.g*., o-180 is *Hans-*LanM, and o-621 is *Mex-*LanM). We expressed this library (SpyTagged) in 96-deepwell block format and individually conjugated variants to CPG:SpyCatcher for SpyCI-LAMBS. [*M*_tot_] is the sum of the metal concentrations across all desorbed ions in the tRE solution (all desorption fraction volumes are the same), which was used as a proxy for the proteins’ expression level. For reference, *Mex*-LanM is a highly-expressed protein with an average [*M*_tot_] = 102 ± 13 ppb, whereas the CPG:SpyCatcher (no LanM) background has an average [*M*_tot_] = 6 ± 2 ppb.

Based on prior work, we expected some orthologs in this library to naturally exhibit poor expression or solubility^19^, and indeed observed a zero-inflated distribution (left-tail kurtosis due to many orthologs having low [*M*_tot_]) comprised of LanMs from the *Bradyrhizobium* and *Methylobacterium* genera (Supplementary Figure 7-Supplementary Figure 9). Nonetheless, we showed earlier (Extended Figure 3g) that 1-10% the lysate quantity of o-621 still generated informative log*D* profiles, so we did not remove datapoints where the average [*M*_tot_] values were below that of the CPG:SpyCatcher control in case they had a meaningful association with phylogeny. By colormapping average [*M*_tot_] values with metadata to assess the randomness within each block (Supplementary Figure 10-Supplementary Figure 12), we found no obvious patterns indicative of human error associated with the handling of individual blocks. Instead, the spread of the [*M*_tot_] values were associated with their phylogeny (Supplementary Figure 13-Supplementary Figure 14), and co-occurrence with RE-dependent enzymes *xoxF* and *exaF* within the host genome (Supplementary Figure 15).

log*D* profiles were then computed from the metal ion concentrations (Supplementary Note 2, Supplementary Figure 16-Supplementary Figure 23). Using uniform manifold approximation and projection (UMAP) and agglomerative hierarchical clustering (Figure 2a), we identified 8 clusters based on manual inspection of the averaged profile per selectivity cluster (Figure 2b), and the silhouette score (Supplementary Figure 24). The eight clusters were labelled Cluster 0 to Cluster 7 (herein notated as “C0” to “C7”). Since C6’s averaged log*D* profile appeared to be a point of divergence in the selectivity of LanM orthologs for the C0**→**C2**→**C3, C5**→**C7 and C1**→**C4 trajectories; and because its taxonomic composition was a blend of C0, C5 and C1’s dominant families (Extended Figure 4, Supplementary Figure 6), we used C6 to anchor our interpretation. C0 members have three partitions: those with sub-background [*M*_tot_] values located between C2 and C3, those with [*M*_tot_] values well above the background located closer to C6, and a *Mex*-LanM satellite (three biological triplicates of o-621). Above C6, C1 and C4 members share the common feature of higher La^III^ selectivity than o-621 and C6, with the most pronounced change in C4 being a Pr^III^ peak characteristic of o-180 when immobilized (in monomeric form). In contrast, C5 and C7 members exhibit dampened selectivity for La^III^. While o-180 from C4 has been shown to improve Nd^III^/Dy ^III^ separation due to its higher LRE selectivity compared to o-621^19^, the C5**→**C7 trajectory appeared to be a unique class of LanMs given the greater intra-LRE selectivity. Critically, C5 and C7 had [*M*_tot_] distributions (Figure 2c) that included members with equivalently high [*M*_tot_] values as o-621, which indicated there were candidates with strong protein expression and separation potential.

**Figure 2.**
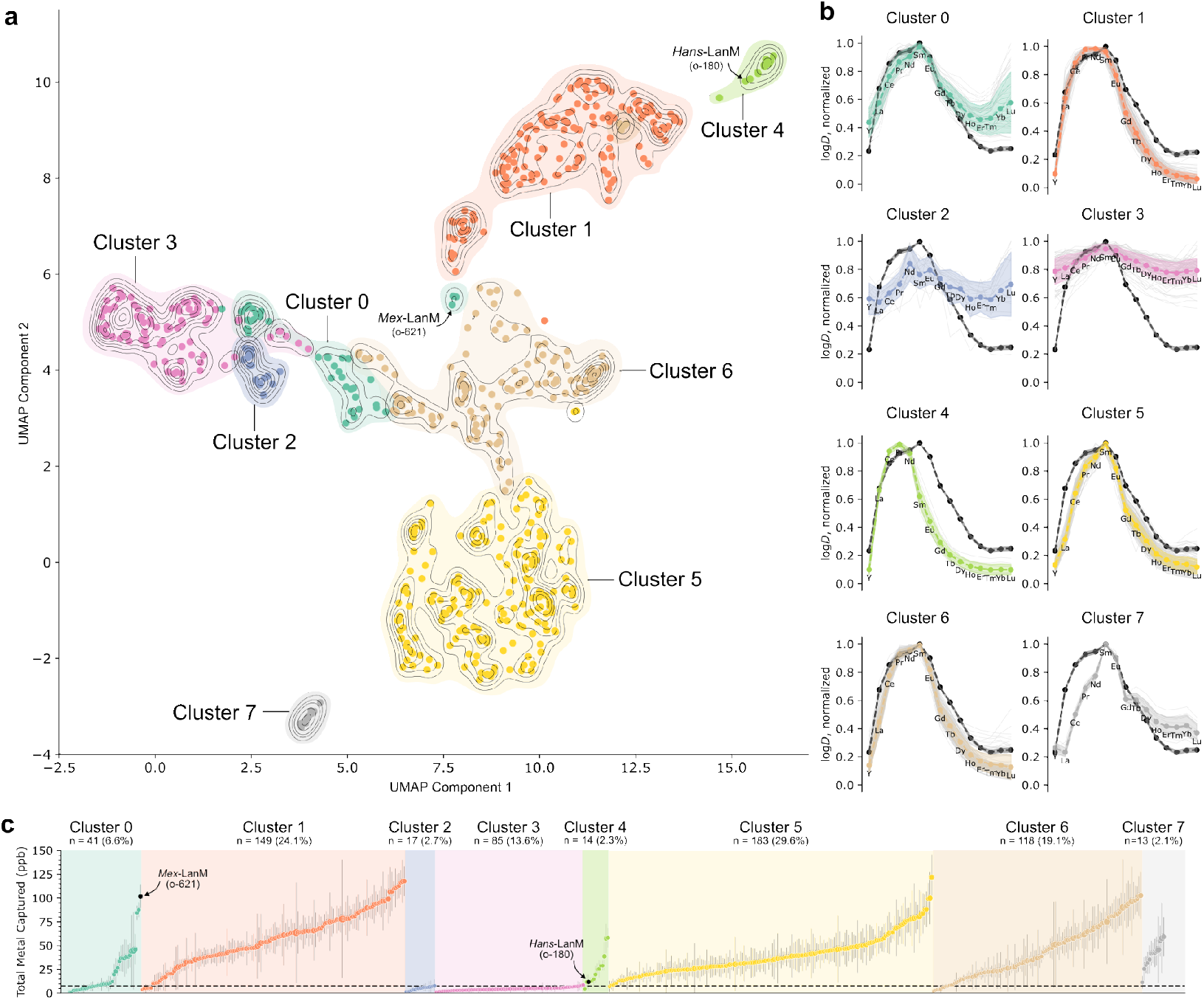
The LanM selectivity landscape. (a) Selectivity clusters. The average of normalized log*D* data (from technical replicates, *viz*. the same CPG:LanM preparations used three times) for all 621 unique LanM orthologs was dimensionally reduced by uniform manifold approximation and projection (UMAP) and clustered using a hierarchical agglomerative approach with contour lines to approximate selectivity cluster boundaries. Each of the eight selectivity clusters (indexed at 0) is assigned a coloring scheme consistent throughout this manuscript. Three biological replicates (*viz*. three different prepartions of the CPG:LanM) of *Mex*-LanM (o-621) and one biological replicate of *Hans*-LanM (o-180) are marked to serve as benchmarks and positive controls (n = 3 technical replicates). (b) Representative normalized log*D* selectivity profiles per cluster. o-621 is plotted in black, with dark grey dashed lines. Individual profiles for each ortholog are shown as trace gray lines. The average of all individual profiles for a particular cluster’s members are shown in color, with a 95% confidence interval error region. (c) Ascending rank order of average total metal captured [*M*_tot_] per ortholog (from the same data as in (a)). Cluster labels and number of members in each cluster are included above the plot. o-621 and o-180 are highlighted as black circles. Error bars represent s.d. (n = 3, technical replicates). A dotted horizontal black line at 6.4 pbb marks the average [*M*_tot_] for non-specific background adsorption of the metal ions to CPG:SpyCatcher.

### Painting a phylogenetic portrait of LanM selectivity

To explore the evolutionary relationship between sequence and function, we prepared an annotated phylogenetic tree of the LanM sequences to produce a family portrait of their selectivity (Figure 3, Supplementary Data 2). Specific nodes are notated as “nX” where “X” is the node identifier number. The tree contained five major branches based on the majority taxonomic composition (herein referred to as “MB”) and revealed strong associations between some clades and the selectivity clusters. Major Branch I (MBI) is comprised of C2 and C3 members from the *Rhizobiaceae* family (n1148), and environmental metagenomes (n1166). MBII was almost entirely comprised of pink-pigmented facultative methylotrophs of the genera *Methylobacterium* and *Methylorubrum*, with some exceptions. MBII also featured strongly supported clades like n1068, n897, and n914 with high homogeneity for C5 selectivity. All C7 members were located under n879, suggesting their log*D* profile could be an evolutionary offshoot of C5. Importantly, the *Hans* cluster that we previously reported on was recapitulated in our tree as MBIII under the strongly supported n1123, and had strong associations with C4 selectivity typical of o-180^19^. Akin to MBII, MBIV was comprised of other families under the *Hyphomicrobiales* order that had high C5 selectivity homogeneity, including *Beijerinckiaceae* (n842, *Methylocella* dominant) and *Methylocystaceae* (n893 – *Methylosinus* dominant, n910 – *Methylocystis* dominant), both families known for methanotrophy. Finally, nearly every member of the genus *Bradyrhizobium* were found in MBV, except for the nine near n896, and distant o-586 in MBI. While MBV clades had weaker bootstrap values than those in other major branches, they typically displayed high homogeneity for C1 selectivity (n631) and C6 selectivity (n775). No C5 selectivity was observed in MBV, which indicated a strong association between the protein sequence of MBV members and C1/C6. This was corroborated by the interspaced and high selectivity homogeneity clades n821/n851 for C6, and n826/n767 for C1, located evolutionarily upstream of n775 and n716. The consistent observation of high selectivity homogeneity across clades that also exhibited high monophyly provided strong evidence to support the hypothesis that RE selectivity of LanM orthologs are evolutionarily linked, while also highlighting SpyCI-LAMBS’ ability to robustly discern selectivity differences between groups of related LanMs.

**Figure 3.**
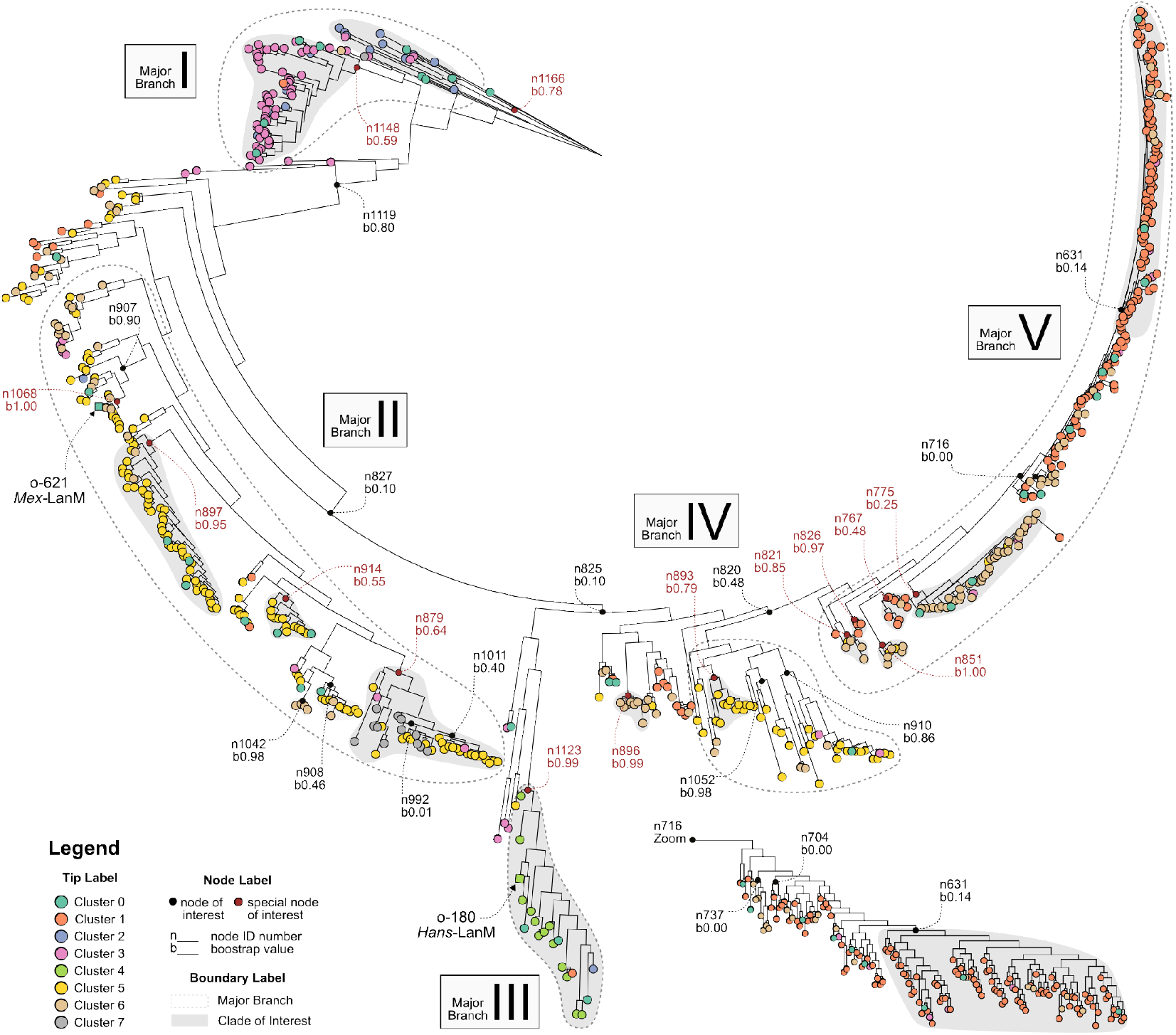
A family portrait of LanM ortholog selectivity. Selectivity cluster data was annotated as circular tip labels in a phylogenetic tree representing 621 unique orthologs, following the same color scheme as Figure 2. Square tip labels for *Mex*-LanM (o-621) and *Hans*-LanM (o-180) are denoted as reference positive controls. Node labels are provided to demark branches of interest (black) and those discussed in the main text (red). Major branches I-V are delineated by a dotted grey perimeter, and clades of interest are delineated by a shaded grey area. A zoomed-in view of the n716 descendants from major branch V is shown separately (bottom right).

### EF-hand sequence features do not solely drive RE selectivity

To contextualize the selectivity with the sequence and structure, we colormapped the MSA conservation scores from the library onto the o-621 crystal structure and the consensus sequence logos of each EF-hand (herein “EF” for brevity) for each selectivity cluster, and focused on EF2 and EF3 thought to have the highest RE affinities (at the time)^17,20^. We observed high levels of conservation (>0.75) for EF residues associated with the first coordination sphere of RE binding, as well as those whose sidechains faced the interior of the protein, likely playing a role in protein folding and stability (Figure 4a). We focused our attention on the EF-hands where the most notable (dis)similarities were associated with the different selectivity clusters (Figure 4b). EF-hand residues at the 1^st^, 3^rd^, 5^th^, 9^th^, and 12^th^ position were generally conserved Asp or Glu; as well as the 6^th^ position Gly. In contrast, residues flanked by these positions (*e.g*., 2^nd^ position) are generally less conserved and have sidechains positioned towards the external environment (Figure 4c).

**Figure 4.**
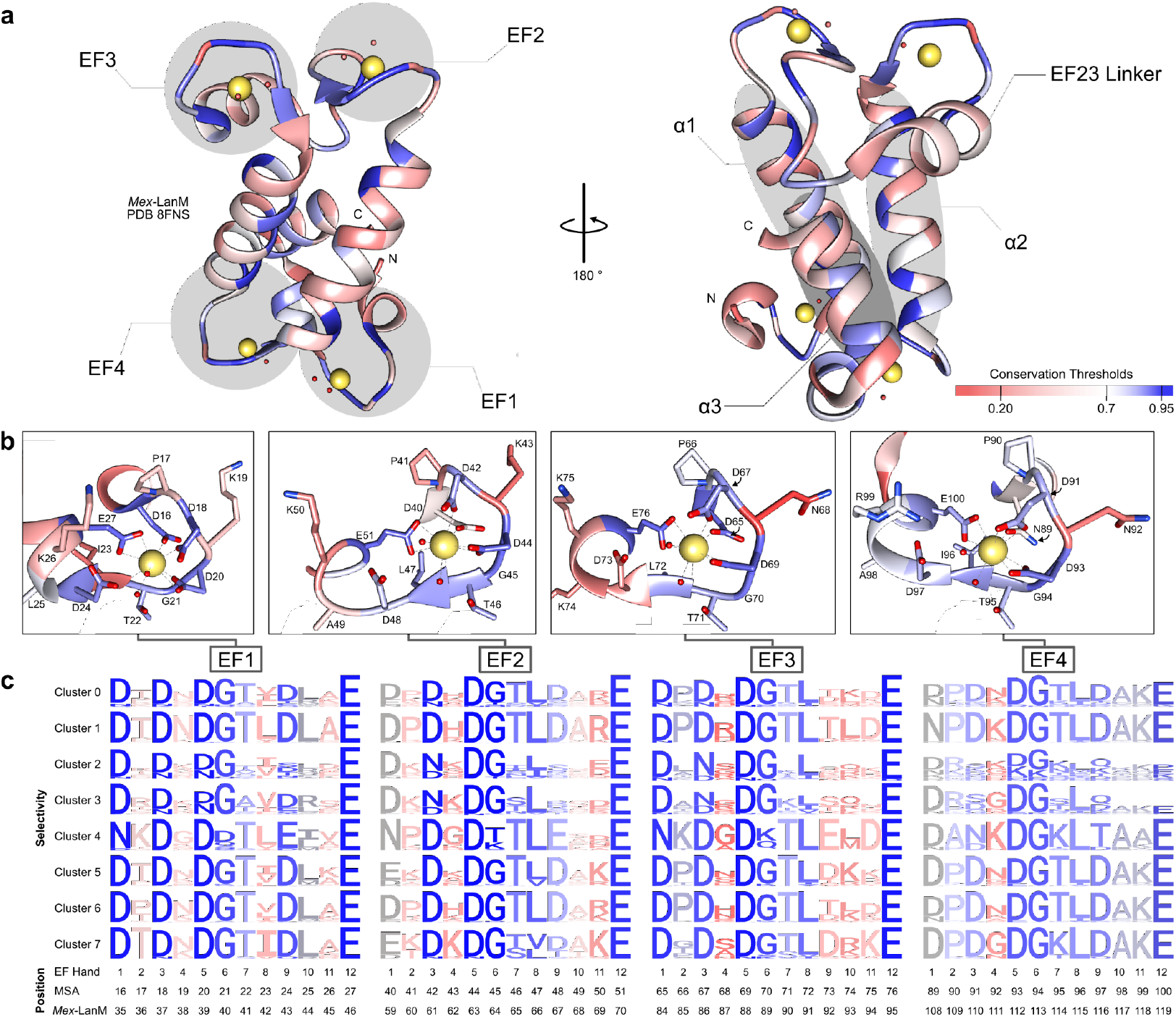
Sequence-structure-selectivity conservation patterns in LanM EF-hand motifs. (a) Left: Front view of *Mex*-LanM (PDB ID: 8FNS)^20^ with EF-hand motifs indicated. Right: Rear view of *Mex*-LanM (o-621) with structural components indicated. Coloring of ribbons indicate conservation score of each position in o-621 with respect to the other LanM orthologs. (b) Zoom-in of EF-hand motifs with residues labelled. Numbering corresponds to the position in the global multiple sequence alignment (MSA). (c) Sequence logos are generated for each selectivity cluster’s EF1, EF2, EF3, and EF4. Coloring follows the same scheme as (a,b). Positional index is provided on the bottom to relate the EF-hand position, the global MSA, and the original o-621 sequence.

There were four exceptions to this trend. First, the majority of C2 and C3 members had no prolines in the 2^nd^ position of any EF, a non-conserved Asn at the 3^rd^ position of EF2 and EF3, weak Thr conservation at the 7^th^ position of EF1-3, as well as a truncated EF4 sequence in orthologs from the *Ensifer, Rhizobium*, and *Sinorhizobium* genera. These features differ from those thought, on the basis of biochemical and structural studies, to confer high selectivity of LanMs for lanthanides and instead are more consistent with traditional Ca(II)-binding EF proteins, which might account for the flat RE selectivity at pH 3 (*vide infra*). Second, C4 members possessed a unique Asn in the 1^st^ position of EF1-3, and the substitution of the conserved Gly at the 6^th^ position with larger residues like Lys or Gln that alter interactions with the loops’ backbones^19^. Third, C1 EF4 possessed a strongly conserved Asn in the 1^st^ position. Interestingly, o-621’s EF4 also possessed an Asn at this same position while other C0 members did not, which we suspect was a contributor to o-621’s log*D* profile being classified into C0 (*viz*. closer to C1) despite its nearest phylogenetic neighbors under n1068 virtually all being C5 or C6. Fourth, C5 and C7 possessed a unique, strongly conserved Glu at the 1^st^ position of EF2, and a more prominent Lys in the 2^nd^ and 4^th^ position of EF2.

We next used SpyCI-LAMBS to determine whether the uniquely conserved amino acid residues within EF2 and EF3 of C4 and C5 LanMs are critical for conferring their distinct intra-RE selectivity profiles (Supplementary Figure 25a). Representative C4 (o-380) and C5 (o-36; *vide infra*) LanMs were selected based on their high [*M*_tot_] values and subjected to a panel of substitutions to make their EFs resemble canonical *Mex*-LanM EFs. For o-36, the Glu and Thr at the 1^st^ and 2^nd^ positions of EF2, as well as the Asp and Thr at the 1^st^ and 2^nd^ positions of EF3, were targeted for substitutions. For o-380, the Asn, Gly, Lys, and Glu residues at the 1^st^, 4^th^, 6^th^, and 9^th^ positions of EF2 and EF3 were targeted for substitutions (Supplementary Figure 25a). All mutants of o-36 retained their higher selectivity against La(III) compared to *Mex*-LanM while all mutants of o-380 retained selectivity peaks around Pr(III) and high LRE vs HRE selectivity relative to Mex-LanM (Supplementary Figure 25b); although, N_1_D and E_9_D mutations in o-380 slightly dampened the LRE vs. HRE selectivity by yielding higher log*D* values for Dy^III^ through Lu^III^. In addition to individual substitutions, we also converted o-36’s EF2 to EF3, EF3 to EF2, or completely swapped their sequences with each other. We saw no significant departure from the o-36 wild-type log*D* profile except for the variant with two copies of the more o621-like EF3, which exhibited a flatter selectivity profile among both the LREs and HREs. These results suggest that the conserved differences in EF motifs between clusters are not the sole drivers of the differing intra-RE selectivity profiles. In light of the differences in [*M*_tot_] between the variants (Supplementary Figure 25c), our results also do not rule out important impacts of these substitutions on protein expression, solubility, binding affinity, kinetics, or (in the case of o-380) dimerization.

### LanM selectivity is consistent on-resin and in-solution

In prior work, we found the relationship between apparent dissociation constants (*K*_d,app_) determined in-solution and log*D* values determined on-resin to agree on the same overall selectivity trends for the reported *Mex*-LanM^6,17^ and *Hans*-LanM^19^. This is important because it indicates the observed selectivities on-resin are not immobilization artefacts. For example, *Hans*-LanM’s log*D* values from SpyCI-LAMBS are consistent with the determined *K*_d,app_ values for the R100K (monomeric) variant (a 27-fold weaker affinity for Dy^III^ compared to Nd^III^)^19^. To investigate whether this notion held true across the selectivity clusters, we purified representative proteins (o-412, C1; o-543, C2; o-585, C3; o-36 and o-127, C5) and studied their binding properties in-solution with several REs. We first confirmed whether they had characteristic binding stoichiometries typical of *Mex*-LanM and *Hans*-LanM by using a standard weak chelator (xylenol orange, XO) as a binding competitor^26^. Purified o-543 (C2) and o-585 (C3) were unable to compete with XO for La^III^-binding (Supplementary Figure 26), corroborating their low [*M*_tot_] values and flat log*D* profiles seen in SpyCI-LAMBS. All other purified LanMs displayed roughly 2.0 binding equivalents (Figure 5), so we next assessed whether they lack pre-ordered secondary structures in their apo states, which would be ideal for label-free direct measurements of their *K*_d,app_ values using circular dichroism (CD) spectroscopy (Supplementary Figure 27). We found o-412 and o-36 had unfolded apo states like *Mex*-LanM (suitable for CD), whereas o-127 presented an unusually pre-ordered structure indicative of α-helices, and thus its *K*_d,app_ was instead determined by spectrofluoremetry.

**Figure 5.**
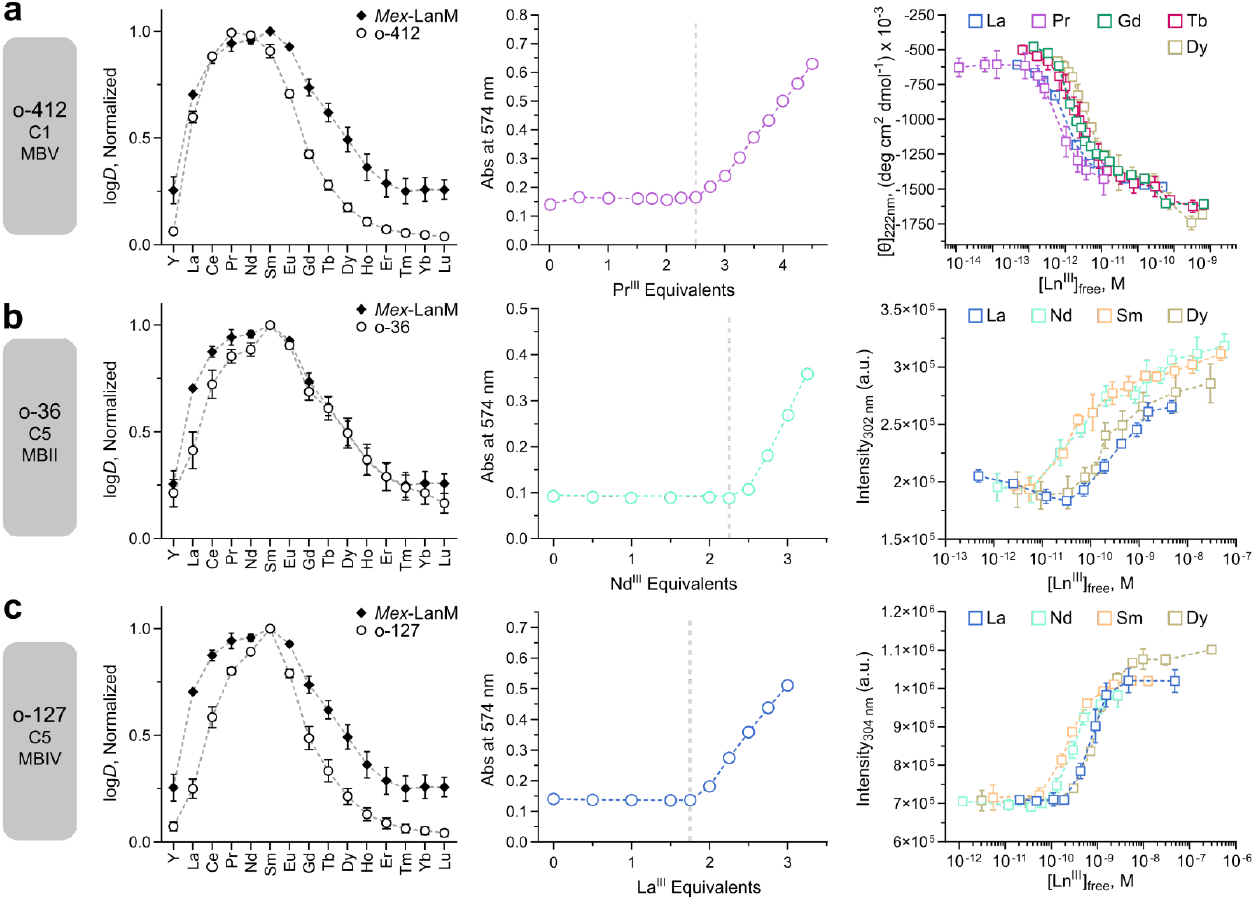
Biochemical characterization of representative orthologs with unique selectivities. (a) o-412. (b) o-36. (c) o-127. Left panels: SpyCI-LAMBS log*D* profiles. Each datapoint represents an average of three biological replicates (*viz*. individual preparations of the CPG:LanM material from different cell lysates). Centre panels: Xylenol orange (XO) plotted against RE molar equivalents added for estimating stoichiometry of each LanM ortholog (n = 1). Right panels: *K*_d,app_ determination at pH 5.0 by (a) circular dichroism spectroscopy (o-412, 10 μM) or (b,c) tyrosine emission (o-36 and o-127, 2 μM each). La^III^ and Pr^III^ were buffered by DTPA, while all other metals were buffered with EGTA. Datapoints represent three independent titrations with s.d. shown, except for o-412 with Gd^III^ (n = 1) and La^III^ (n = 2).

Based on o-412’s log*D* profile compared to *Mex*-LanM (Supplementary Figure 17), we expected similar relative affinities among LREs and marginally weaker relative affinity for HREs (Supplementary Table 2). Using CD, we indeed found o-412 had similar relative affinities for the LREs (1.6-fold difference between Pr^III^/La^III^), and weaker relative affinity for HREs (6.2-fold difference between Pr^III/^Dy^III^ for o-412 compared to 4.7-fold difference between the similarly distant (among the Ln^III^ series) Nd^III^/Ho^III^ for *Mex*-LanM^17^. The C5 members o-36 and o-127 were of particular interest, due to their steeper selectivity against La^III^ according to SpyCI-LAMBS. We used spectrofluorometry to allow direct comparison of these proteins. Both LanMs displayed notably weaker relative affinity for La^III^ than that observed for *Mex*-LanM and the monomeric version of *Hans*-LanM (R100K) (Figure 5b,c, Supplementary Table 2); o-36 had a striking 8.7-fold difference in affinity between Nd^III^/La^III^, whereas this difference was 3.5-fold for o-127 and 1.7-fold for *Mex*-LanM. This result indicates that the phylogeny of o-36 (MBII, primarily *Methylobacterium*) likely represents evolutionarily distinct LanMs that have steeper LRE selectivities compared to those from MBIV that o-127 represents (primarily *Methylosinus, Methylosinus*, and *Methylocella*). A LanM from C7 (o-21), which also displayed steep selectivity against La^III^ in SpyCI-LAMBS expressed poorly and could not be purified in sufficient quantities for further characterization. Taken together, these data suggest that the selectivity trends determined by SpyCI-LAMBs (and observed in SpECS, *vide infra*) are reflective of the behavior of the in-solution proteins. Additional considerations and caveats are discussed below.

### C5 LanMs reject lanthanum for enhanced RE separations

To identify LanM orthologs with better intra-LRE separation than *Mex*-LanM (o-621) using a pH-only desorption scheme, we developed SpECS for rapid screening of a protein’s separation potential (Figure 6a). From each cluster, we initially chose members with the highest [*M*_tot_] values (Figure 2c). We then picked the final five candidates based on their high soluble protein expression levels and unique separation factor (SF) heatmaps (Supplementary Figure 29 to Supplementary Figure 34): o-25 (C7) from *Methylobacterium aerolatum*, o-36 (C5) from *Methylobacterium sp*.13MFTsu3.1M2, o-40 (C5) from *Methylobacterium sp*.P1-11, o-257 (C1) from *Bradyrhizobium japonicum*, and o-380 (C4) from *Methylopila sp*.73B (Extended Figure 5, Supplementary Figure 35). Each candidate was conjugated to CPG:SpyCatcher and packed into a narrow tubing yielding an aspect ratio (AR) of 118 (Figure 6a). These capillary-like columns were then installed on a modified pump-fractionator apparatus (Supplementary Figure 36) to perform column experiments simultaneously.

**Figure 6.**
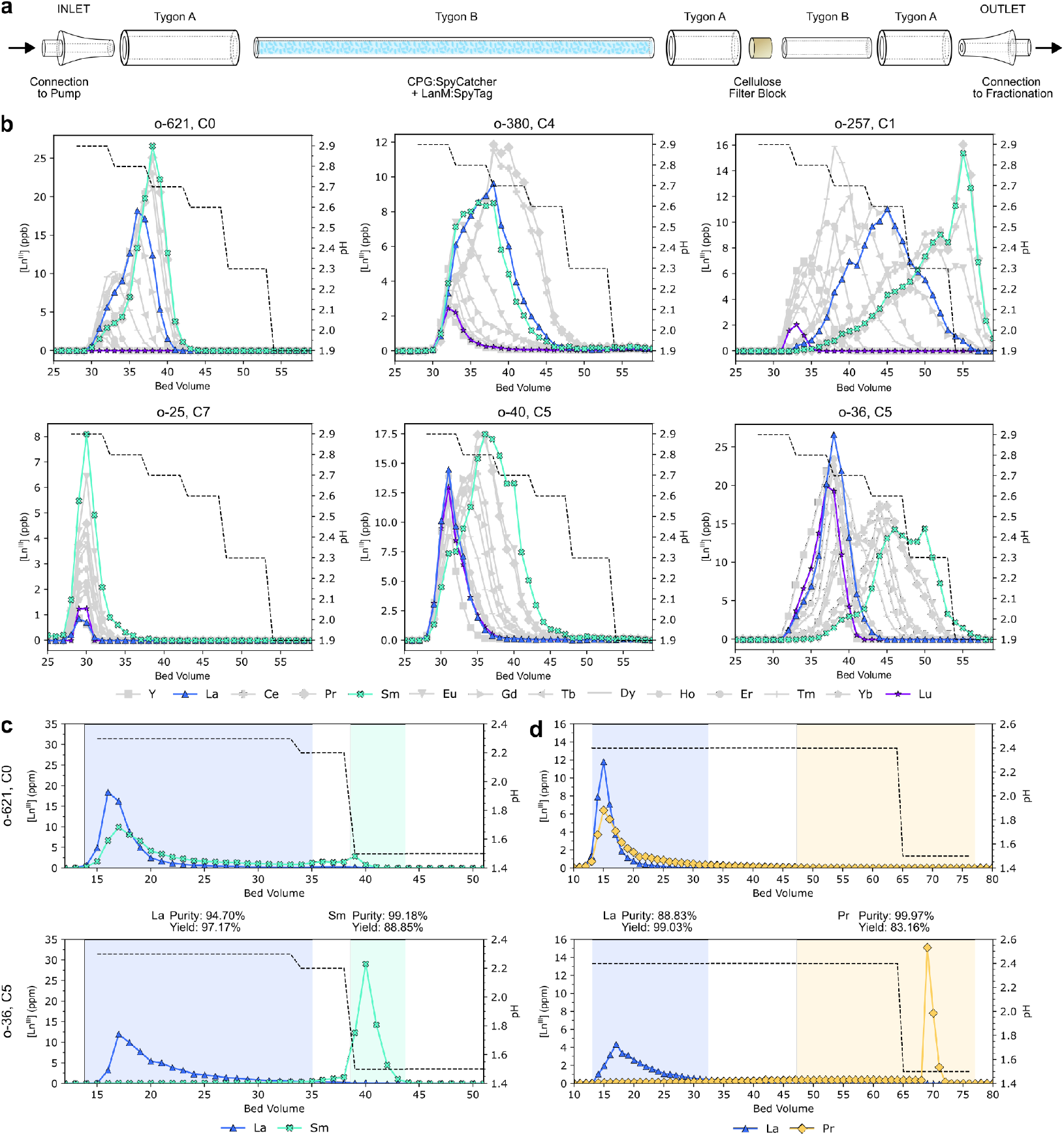
Column testing of top candidates. (a) Assembly of high aspect ratio SpeCS columns for higher throughput separation tests using common liquid chromatography components. (b) Stepwise pH-based desorption using increasing [HCl] steps (black dotted line). tRE-loaded SpeCS columns contained LanM orthologs of interest immobilized via SpyTag-Catcher chemistry directly from cell lysate. 1 bed volume = 0.1 mL, corresponding to the total volume occupied by the CPG:LanM material inside the SpeCS column space. La^III^, Sm^III^, and Lu^III^ are highlighted in color to demark differences in the La-Sm plateau experienced by the different orthologs (blue and cyan, respectively), as well as the theoretical weakest affinity RE (Lu^III^, purple) to provide a sense of differences in affinity. All other REs are shown as grey traces with unique markers per RE. n = 1. (c,d) Traditional column separation performance of o-621 (top panels) and o-36 (bottom panels) with binary solutions of La^III^ and Sm^III^ (left) or Pr^III^ (right). A simplified stepwise pH-based desorption using small [HCl] increments (black dotted line) was used to purify the non-La^III^ RE. Shaded regions demark the regions (a range between bed volumes, BVs) used to compute the purity and yield of La^III^ and Sm^III^/Pr^III^ after completing the run. 1 BV was 0.8 mL. n = 1.

All five candidates generated unique desorption chromatograms compared to o-621 (Figure 6b, Extended Figure 6). The REs eluted from the o-621 (C0) benchmark between pH 2.9-2.7 across 13 bed volumes (BVs) in the order of its log*D* selectivity (Figure 6b), with no Lu^III^ able to bind. Like o-180^19^, o-380 (C4) had peak selectivity for Ce^III^ and Pr^III^ based on SpyCI-LAMBS, which was reflected in the order of o-380’s RE elutions and the total area under the curves per RE (*viz*. its capacity). REs eluted from o-257 (C1) starting at pH 2.9 and did not complete desorption by the end of the pH 1.9 step (28 BVs), suggestive of high affinity. We expected o-25 (C7) to present the best separation of La^III^ and Sm^III^ based on its log*D* profile (Extended Figure 5), but we observed immediate desorption of all REs at the single pH 2.9 step (5 BVs) suggestive of weak REE binding affinity or poor expression similar to that observed for o-27 (C7). While both o-40 and o-36 displayed C5 selectivity in SpyCI-LAMBS, o-36 had larger capacities per RE overall (matching o-621) and strinkingly higher-resolution separation of La^III^ and Sm^III^, compared to o-40. Since SpECS columns were operated at three orders of magnitude lower capacities than traditional LanM-agarose columns (up-to 100 ppm/mL_agarose_ capacity), we anticipated loading purified o-36 to higher densities on agarose columns could significantly boost its separation potential.

We next purified o-36 and o-621 with a -GSGC tag to fabricate 0.8-mL agarose columns with a confirmed 2 μmol_RE_/mL_agarose_ capacity, determined by Sm^III^ breakthrough (Supplementary Figure 39). o-36’s log*D* profile determined using the traditional column approach was nearly identical to the profile determined using SpyCI-LAMBS, especially in the La^III^-Nd^III^ region (Supplementary Figure 38). Both columns were fed a binary 50:50 solution of La^III^ and Sm^III^ with bound REEs eluted using a 2-step pH gradient (Figure 6c,d, Extended Figure 7, Supplementary Figure 40). o-36 achieved 94.7 mol% La purity at 97.2% yield, and 99.2 mol% Sm purity at 88.8% yield, whereas o-621 could not achieve separation of these two REs under the same conditions. Motivated by this result, we tested the more challenging Pr^III^/La^III^ separation using a 50:50 binary solution of the two REs. With a single step-drop from pH 2.4 to 1.5, o-36 achieved >99.9 mol% Pr^III^ with 83.2% yield. To confirm this ability to reject La^III^ was reproducible, we re-fabricated new 0.4-mL agarose columns with a new batch of proteins. Without optimization of the elution timings and pH increments to accommodate the smaller scale, we still achieved a 98.6 ± 1.1 mol% Sm^III^ purity with a 72.6 ± 1.1% yield, and 96.5 ± 0.2 mol% Pr^III^ purity with a 72.0 ± 5.8% yield (Extended Figure 8). In light of this demonstration, we name o-36 as “*Melba*-LanM”, a prototypical representative of the C5 selectivity cluster that reproducibly overcomes the LRE selectivity plateau of *Mex*-LanM and *Hans*-LanM.

## Discussion

In this work, we developed a technique to measure RE selectivity *en masse*, and applied it to 621 LanM orthologs. SpyCI-LAMBS generated reproducible log*D* profiles that matched published work^6,19^, and was robust to several procedural alterations or experimental fluctuations, including variations in protein concentration. Eight selectivity clusters were identified, and were strongly associated with the LanM orthologs’ phylogeny. This included C5, predominately from *Methylobacterium*, which presented higher intra-RE selectivity within La^III^-Nd^III^ compared to *Mex-*LanM and other orthologs, enabling reproducible high-purity separation of La^III^/Sm^III^ and La^III^/Pr^III^ binary solutions.

These results for *Melba-*LanM are significant since La^III^ and Ce^III^ are valued around $1/kg (at the time of writing), yet comprise up to 75% of the tRE distribution in feedstocks such as bastnasite.^11^ As such, the ability to selectively enrich for only the LREs critical for magnet manufacturing (*e.g*., Pr^III^, Nd^III^, Sm^III^) while rejecting La^III^ would improve RE separation process economics. Importantly, this separation can be accomplished without the use of an exogenous ligand (*e.g*., citrate) as required previously for Nd^III^/Pr^III^ group separation from La^III^/Ce^III^ by *Mex*-LanM^11^. This enables facile conversion of the separated REs into a form amenable to downstream processing (*e.g*., oxides). While the LRE separation factors (SF) of *Melba*-LanM are comparable with those for a recently discovered and further engineered protein, *Mex*-LanD– E75Q/E78A (SF_Pr/La_ of 5.1), which enriches for Nd^III^/Pr^III^ over La^III^ through selective dimerization^27^, the compatibility of *Melba-*LanM with a column-based process enables the propagation of its separation factors over multiple theoretical plates, resulting in effective intra-LRE ion separation. Additional work would be required to leverage the dimerization mechanism of LanD on-column to facilitate a direct comparison of separation performance. In other recent work, HEW5 from *Nocardioides zeae* was shown to achieve a 50:50 binary separation of a La^III^/Nd^III^ solution with >90% purity and 90% yield^16^. While SFs were not determined for immobilized HEW5 to enable quantitative comparison of on-column selectivity, the *K*_d,app_ values determined for free and immobilized HEW5 for La^III^ and Nd^III^ suggest shallower LRE selectivity compared to *Melba*-LanM. We suspect this could explain its less resolved LRE separation despite deploying a larger column (0.4 mL for *Melba*-LanM compared to 7 mL for HEW5) and a more complex desorption regime (single-step pH drop for *Melba*-LanM compared to a pH gradient for HEW5).

A hallmark of the lanmodulin family is preferential binding to LREs (namely, Ce^III^-Sm^III^) over HREs, as reinforced by our comprehensive selectivity profiling. This property has enabled industrially relevant separations, with *Hans*-LanM being the most effective ortholog characterized to date^19^. Our data confirm that the Hans cluster^19^ (MBIII, C4-dominant) includes the most LRE-selective LanMs with a selectivity maximum shifted the furthest toward the LREs (peaking at Pr^III^). However, this study has unearthed several orthologs with similarly steep HRE rejection, approaching that of the C4 orthologs (at least when immobilized as monomers). For example, C1 orthologs like o-412 (from *Methyloligella* sp.GL2) have selectivity maxima centered around Nd^III^ and greater selectivity against Dy^III^ than *Mex*-LanM. When immobilized onto agarose, o-412 had a SF_Nd/Dy_ of 10.5 (Supplementary Table 3) compared to the reported 13.7 for *Hans*-LanM (R100K)^19^. In addition to having high intra-LRE selectivity, some C5 members (from MBIV) have similar or even higher Nd^III^/Dy^III^ selectivity compared to *Mex*-LanM (Supplementary Figure 21), which we confirmed through biochemical characterization of o-127. From a process perspective, these C5 subcluster orthologs may facilitate a broader range of RE separations given their steep intra-LRE and intra-HRE selectivities. From a design perspective, the protein scaffolds for C4 and C5 orthologs may therefore represent two different “trajectories” in protein sequence and/or structural space for re-designed LanMs with improved selectivities.

The mutational analysis of EF2 and EF3 from *Melba*-LanM and o-380 highlights the complexity of the principles that govern the lanmodulin family’s differing intra-LRE selectivities. While C4 and C5 members contained uniquely conserved residues within their EF hand domains, their targeted substitution yielded minimal change to the log*D* profiles. We argue the origins of intra-RE selectivity across the tested 621 members of this protein family likely arises from a delicate and deliberate interplay between the EF-hands and the rest of the structure. This interplay could be decomposed into residue-residue interactions within 1) the EF-hands, 2) between them (*e.g*., the linker between EF2 and EF3), 3) within the scaffold (comprised of the α-helical bundle core), and 4) between the scaffold and the EF-hands. It is known that *Mex*-LanM deploys a nine-coordinate geometry to bind to Nd^III^ (PDB: 8FNS), whereas *Hans*-LanM uses a ten-coordinate geometry for La^III^ and nine-coordinate for Dy^III^ (PDB: 8DQ2). While an x-ray structure for *Melba*-LanM has yet to be solved, we suspect C5 members have evolved a divergent arrangement of these interactions to transition between the high (10) to lower (9 or perhaps even 8) coordination numbers across a narrower range of ionic radii^28^, differently from *Mex*-LanM and *Hans*-LanM. Based on our brief mutational study of *Melba*-LanM, we suspect its La^III^ rejection is rooted in how its scaffold shapes the secondary coordination sphere (and beyond).

Several published works suggest that EF-hands are not the sole drivers of selectivity despite directly coordinating with the RE ions. Structural characterization of the RE-bound *Hans*-LanM revealed a more-extensive secondary sphere, hydrogen-bonding network that positions the coordinating EF-hand residues, thus increasing sensitivity to RE ionic radius^19^. Similarly, molecular dynamics simulations for *Mex*-LanM indicated that an interaction network involving EF2, EF3, and the intervening domain connecting these two yielded higher accuracy energy scores when compared to a network that only accounted for interactions within the EFs^29^. Additionally, through substitutions of Asp in the 9^th^ positions of *Mex-*LanM’s EFs, which forms a hydrogen bond with a metal-coordinating water, Mattocks *et al*. (2022) showed that secondary sphere interactions can tune actinide vs. lanthanide selectivity^22^. Overall, our mechanistic understanding of intra-RE selectivity in LanMs remains limited, underscoring the need for further biochemical and biophysical studies across a broader range of LanMs.

The close correspondence between phylogeny and selectivity reported in this work supports an evolutionary basis for the distinct LanM selectivity clusters, which we suspect are tied to the environments from which these microorganisms originate. For example, C1 and C5 members represent the dominant selectivity clusters from MBV and MBII/MBIV, respectively. The former include *Bradyrhizobium* species typically found as root endophytes, whereas the latter include *Methylobacterium, Methylosinus*, and *Methylocystis* that are mostly found in the bulk soil environment, rhizosphere, or phyllosphere^30,31^. These differences in preferred habitat or mode of interaction with plants could subject these organisms to different RE distributions and speciation that impact RE bioavailability. Larrinaga *et al*. (2024) proposed a role for *Mex*-LanM in sequestering catalytically disfavored metals like Nd^III^ and Sm^III^, in conjunction with *Mex*-LanD, helping to ensure that the catalytically more favored La^III^ and Ce^III^ remain available for metalation of the methanol dehydrogenase XoxF^27^, a mechanism that is harmonious with the involvement of LanM in membrane-vesicle mediated efflux of REs in *M. aquaticum*^32^. LanM and potentially LanD may have evolved selectivity trends that reflect the relative RE abundance in these different environments (and the selectivity of RE uptake). While studies to date have revealed important differences in lanthanide utilization clusters (gene composition and regulation), uptake, and storage in LanM-bearing methylotrophs, the selectivity of this metabolic machinery across diverse environments and RE bioavailability/speciation in these environments remain inadequately characterized. Our work identifies microorganisms where searches for better selectivities might be most profitable.

Finally, SpyCI-LAMBS log*D* profiles are snapshots of LanM’s selectivity under specific conditions (equimolar tRE buffered at pH 3.0) and may not necessarily reflect their physiological selectivity (*e.g*., due to oligomerization)^19,27^ or behavior in complex feedstocks. Seidel *et al*. (2024) showed the separation factors for RE ion pairs with *Mex*-LanM can vary widely based on the ratio of RE concentrations present, on account of mixed RE complex formation by its EF-hands^18^. Since hetero-ion complexation tends to favor complexes of hetero-ion pairs with dissimilar radii than the homo-ion complex with the same radii, the selectivities reported here likely underestimate the LanMs’ true selectivities. As such, we anticipate log*D* profiles will be influenced by the relative RE compositions in real feedstock solutions, warranting SpyCI-LAMBS under a broader range of conditions to comprehensively inform the RE separation potential of a particular LanM. In future work, we plan to modify SpyCI-LAMBS to study the selectivity (and affinity, by chelator competition) of engineered LanM variants and other metalloproteins with a larger panel of metal compositions. Moreover, laboratory automation has advanced to a point where SpyCI-LAMBS can be performed robotically. This would enable larger screening campaigns to generate the high-fidelity datasets needed to train machine learning models to assist with protein design for LanMs and other metalloproteins for critical metal recovery. The predicted rise in the demand for critical materials will only spotlight the need for more sustainable approaches to mining.

## Acknowledgements

This work was funded by the DARPA Environmental Microbes as a Bioengineering Resource (EMBER) program (Y.J.), and the work at LLNL was performed under the auspices of the U.S. Department of Energy by Lawrence Livermore National Laboratory under Contract DEAC52-07NA27344 (LLNL-JRNL-871871). Distribution Statement A. Approved for public release: distribution is unlimited. The views, opinions, and/or findings expressed are those of the authors and should not be interpreted as representing the official views or policies of the Department of Defense or the U.S. Government. We thank Andrew Gross for ICP-MS technical support, Max I. Goldman for ICP-MS access, and Bentley Lim for turbo colony-picking support. We also thank several colleagues for helpful discussions (Mimi C. Yung, Bentley Lim, Sean P. Leonard, Ziwei Zhong, Chenling Xu, Daniel J. Wackelin, Jennifer L. Chlebeck, Xun Wang, and Wei Li).

## Author contributions

D.M.P conceived and directed the study, with the support of Y.J. P.D. helped conceive the study, cloned the LanM library, developed and deployed SpyCI-LAMBS, carried out the metal separations and ICP-MS measurements; and performed the bioinformatics and data analysis. C.S.M. supported cloning of the LanM library and data analysis. W.C. purified o-36, o-127, and o-412 and performed biochemical characterization with input from J.A.C. Z.D. supported protein immobilizations and column experiments. C.S.K.-Y. supported the bioinformatics analysis. D.M.P. and J.A.S. supported data analysis. S.A.E. purified o-543 and o-585 and performed biochemical characterization with input from J.A.C. P.D. and D.M.P. wrote the paper with input from all authors. J.A.C. supported the preparation of the final manuscript. All authors edited and approved the final manuscript.

